# CAR T-cells dysfunction in the central nervous system is mediated by BBB-induced activation-induced cell death

**DOI:** 10.1101/2025.10.21.683607

**Authors:** Yarden Aharony-Tevet, Orly Ravid, Asaf David Yanir, Amilia Meir, Hadar Levi, Daniel Burstein, Aviya Amram, Tamar Feuerstein, Shai Izraeli, Itzik Cooper, Elad Jacoby

## Abstract

Chimeric Antigen Receptor (CAR) T-cells can eliminate leukemia and lymphoma within the central nervous system (CNS). Following a case with CD19+ relapse in the CNS in the presence of persisting Tisagenlecleucel, we generated several models to study CAR T-cells in the CNS. In both immunocompetent and humanized leukemia-bearing mice, CNS-residing CAR T-cells were shown to have inferior *ex-vivo* killing capacities compared to CAR T-cells harvested from the spleen or bone marrow of receptive mice. In a mechanistic work-up, we identified two contributing factors: First, CAR T-cells cultured in cerebrospinal fluid showed impaired cytotoxic and proliferative function compared to those in serum or other media. Second, using a unique *in vitro* human blood-brain barrier (BBB) model composed of endothelial cells and brain pericytes, we demonstrated that CAR T-cells that crossed the BBB exhibited antigen-independent activation and diminished cytolytic capacity. This led to increased activation-induced cell death of the BBB-migrating CAR T-cells and was confirmed to be independent of the CAR target and the costimulatory domain. Our findings add possible mechanisms of CAR T-cell resistance in the CNS, relevant for CNS hematologic and non-hematologic tumors.

## Introduction

The central nervous system (CNS) is considered a pharmacologic sanctuary site, posing a significant challenge in the treatment of hematologic malignancies with CNS involvement or of other tumors of the brain^1^. This sanctuary status is largely attributed to the blood–brain barrier (BBB), a selective and tightly regulated interface that restricts the entry of many systemically administered therapies, including most chemotherapeutic agents and monoclonal antibodies. As a result, CNS-directed therapy is required in patients with acute lymphoblastic leukemia (ALL) or certain subtypes of lymphoma, and administered via intrathecal route or radiation^2,3^. The incidence of CNS involvement is increased in relapsed ALL^4^. Patients with CNS lymphoma often experience poorer outcomes due to the limited efficacy of conventional treatments in this compartment^5^.

The need for therapeutic modalities that can bypass the BBB has spurred interest in immune-based approaches, including chimeric antigen receptor (CAR) T-cell therapy. CAR T-cells have demonstrated remarkable efficacy in relapsed or refractory B-cell malignancies, particularly ALL and lymphoma^6^. Initial concerns about potential neurotoxicity led to hesitation in using CAR T-cells in patients with active CNS disease. Later emerging clinical data suggest that CAR T-cells are capable of trafficking to the CNS and mediating potent antitumor activity^7–10^.

However, durable remissions following Tisagenlecleucel for isolated CNS relapse of ALL were limited in absence of further therapy^8^. In secondary CNS lymphoma treated with CAR T-cells, although initial remission rates are comparable to real world data of lymphoma without CNS involvement, durability of the response may also be limited^10,11^. This raises concern regarding activity of CAR T-cells in the CNS.

In addition, early clinical trials administering various CAR-T products for primary CNS malignancies reported minimal activity when T-cells are administered intravenously, and have led to repeated CAR-T cell infusions directly to the cerebral-spinal fluid (CSF)^12–14^.

In this work we utilized *in vivo* and *in vitro* models to characterize the phenotype and functionality of CNS-infiltrating CAR T-cells. We demonstrate poor functionality of CNS-residing CAR T-cells, related to the low nutritional status of the CSF and a unique mechanisms of CAR-T cell activation-induced cell death (AICD) triggered by the BBB.

## Materials and Methods

### Ethical approval

All animal experiments were approved by Institutional Ethical Review Process Committees and were performed under Israel Institutional Animal care and use committee approval (1308/21). Human CSF and Serum samples from CD19 CAR T-cell treated patients were collected under an IRB-approved protocol (NCT02772198). Samples were collected on day 30 post CAR treatment while in remission. Serum was separated from blood by 10 min centrifugation in 400 g and supernatant of CSF samples were collected. Samples were stored in −80°C for further use.

### CAR T-cells production

CAR-T cell production was performed as previously described^15,16^. Briefly, human peripheral-blood mononuclear cells were activated using 50ng/mL OKT3, and transduced with an MSGV CAR retrovirus, containing either a CD19-targeting CAR (FMC63-CD28-CD3ζ) or a CD22-targeting CAR (m971-41BB-CD3ζ). CAR-T cells were then grown for 7–11 days. For murine CAR-T production, C57Bl/6 splenocytes were activated using magnetic CD3/CD28 beads at a 2:1 bead-to-cell ratio, and transduced with a MSGV-CD19-targeting CAR retrovirus (1D3-2MZ-CD28-CD3ζ-IRES-GFP, backbone kindly provided by Dr. James Kochenderfer). Murine CAR T-cells were cultured for 5-7 days.

### Cell culture and cell lines

Nalm6 cells were purchased from ATCC. The generation of Nalm6^CD19-KO^ was previously reported^17^. The E2a::PBX murine leukemia cell line was kindly provided by Dr. Terry Fry and Dr. Naomi Taylor. Human cells were grown in RPMI with 10% serum. Murine cells were grown in complete mouse media^18^. For CAR T-cell generation, IL-2 was supplemented in the cultured at a concentration of 100IU/ml (human) or 50IU/ml (murine). Co-culture experiments were performed without additional IL-2.

### *In vivo* models

NOD−SCID−IL-2Rγ− (NSG) and C57BL/6J (B6) mice were purchased from the Jackson laboratories. Six to ten-week-old female mice were tail-vein injected with 1×10^6^ Nalm6 or E2a::PBX cells, and after 7 days treated IV with 5×10^6^ CAR T-cells or untransduced T-cells. Mice were sacrificed 7 days (unless stated otherwise) after CAR T-cells injection and cells from bone marrow, spleen and meninges (CNS) were collected and further analyzed. For long term survival experiments, mice were monitored at least twice weekly for signs of leukemia, graft-vs-host disease, and survival. Moribund mice were sacrificed per institutional guidelines.

### Flow Cytometry and Cell Sort

Flow cytometry was performed on a Cytek Aurora. The antibodies used for cell staining are reported in the supplemental methods. Cell sorting was performed on Sony flow-based sorter or MACS cell separator with LS columns with anti-human CD8 beads and anti-PE beads (Miltenyi). Patient flow cytometry was performed using FACS Canto (BD). CSF was stained with LC-12 (Cytognos) or separate antibodies (see supplemental Table 1). Analysis was done using the FlowJo analysis software.

### BBB in-vitro model

Primary bovine brain pericytes are seeded on gelatin-coated plates while human CD34+-derived endothelial cells are seeded onto Matrigel-coated Transwell inserts. Cells are grown in co-culture for 6 days and acquire BBB properties^19,20^. The top compartment represents and corresponds in its biological properties to the luminal side of the blood vessel and the bottom represents the abluminal brain side. 5×10^5^ CAR-T cells were introduced into the luminal side, and allowed to migrate for 4 hours.

### IFNγ secretion

Cell free supernatant from co-cultures were collected and stored at −20°C prior to analysis. ELISA was performed using human or mouse IFNγ ELISA kit (Invitrogen) according to manufacturer instructions.

### qPCR for CAR construct

CSF of patient was obtained and DNA was extracted with DNA purification kit (Qiagen). To quantify CAR construct expression, TaqMan (PCRBiosystems) based RT-PCR in a Step One Plus system (#8024, Applied Biosystems) was used. The following PCR-program was performed: 20s 95 °C; 95 °C for 15 s, 60 °C for 30 s, repeated 40 times (amplification); 95 °C for 15 s, 60 °C for 1 min, 95 °C for 15 s. The PCR results were evaluated using the Applied Biosystems software. Primers used: CAR CD19-41BB FW: ‘TGCCGATTTCCAGAAGAAGAAGAAG’, CAR CD19-41BB Rev: ‘GCGCTCCTGCTGAACTTC’, CAR CD19-41BB Probe: ‘6-FAM-ACTCTCAGTTCACATCCTC-MGB-BHQ1’, CDKN1a FW: ‘GAAAGCTGACTGCCCCTATTTG’, CDKN1a Rev: ‘GAGAGGAAGTGCTGGGAACAAT, CDKN1a Probe: ‘6-Fam-CTCCCCAGT CTCTTT’

### Statistical analysis

All statistical analysis was done on GraphPad Prism 9 software. Where applicable, non-parametric (Mann–Whitney/Kruskal-Wallis) test or ANOVA were used to obtain p-values of significance across conditions. Survival was analyzed by the Kaplan-Meier method.

## Results

We report on a 16-year-old patient with Ph+ ALL, who presented with an isolated CNS first relapse of her disease 1.8 years after allogeneic HSCT in CR1, when she was off imatinib (**Fig.1A**). In flow cytometry from the CSF, blasts had dim CD45 expression, and were positive for CD19 and CD10 (**Fig.1B)**. The patient was treated with Tisagenlecleucel after bridging with dasatinib and intrathecal chemotherapy, and had prolonged remission with ongoing peripheral B-cell aplasia (BCA). Sixteen months post tisagenlecleucel, the patient had a 2^nd^ isolated CNS relapse, with blasts still expressing CD19 **(Fig.1C)**. Of note CD3+ cells were present in the CSF, PCR was positive for CAR transgene expression (**Fig.1D)**, and peripheral BCA was ongoing (**Fig.1E)**. The patient was treated with intrathecal chemotherapy, steroids and radiation, with clearance of CNS blasts but a 3^rd^ CNS relapse 6 months later was treated with ponatinib. The presence of ongoing peripheral B-cell aplasia, a CD19+ relapse in the CNS, and ongoing CAR-T cell circulating in the CNS, suggest functional impairment of the CAR T-cells in this compartment.

**Figure 1:**
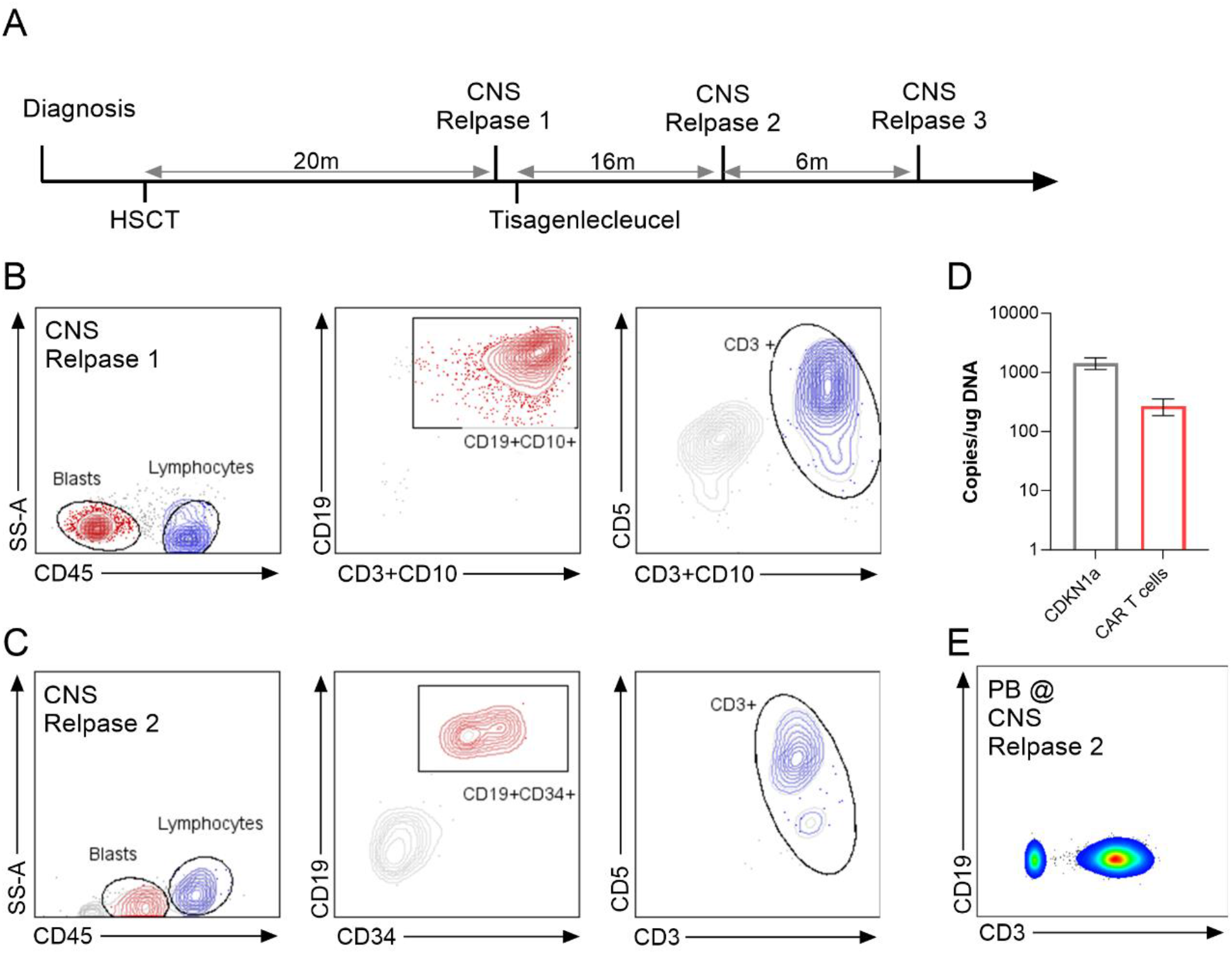
CNS-confined CD19-positive relapse in the presence of CAR T cells and peripheral B-cell aplasia. **A** Clinical course of a patient with multiple central nervous system (CNS) relapses in which samples were analyzed in panels B-E. **B** Flow cytometry plots of cerebral spinal fluid (CSF) at time of CNS relapse 1, showing CD19+ blasts and CD3+ T-cells. **C** Flow cytometry plots of the CSF at time of CNS relapse 2 (16 months post Tisagenlecleucel), showing CD19+ blasts and CD3+CD5+ T-cells. **D** RT PCR for CD19 CAR and CDKN1a in the cell pellet from the CSF at time of CNS relapse 2, normalized to copies per microgram of DNA. **E** Flow cytometry plots of lymphocytes (CD45-high) in the peripheral blood (PB) at time of CNS relapse 2, showing CD3+ T-cells and lack of CD19+ B-cells.

To evaluate CAR-T cell function in the CNS we utilized an immunocompetent model^18^ of C57Bl/6 mice receiving E2a::PBX1 leukemia (based om the TCF3::PBX1 translocation) on day −7, lymphodepletion with cyclophosphamide on day −1 and murine CD19 CAR T cells on day 0 (**Fig. 2a**). We cloned the previously published murine CD19 CAR to include GFP expression after IRES sequence (**supplementary Fig. 1**), and confirmed *in vitro* and *in vivo* efficacy of this construct (**supplementary Fig 2**). *In vivo* CAR-T kinetics demonstrated peak expansion on day 7, as assessed in the spleen, bone marrow (BM) and the CNS of the mice (**Fig. 2b**), along with elimination of CD19+ cells (**Fig. 2C-E**). When assessed at a later time point (day 11), we observed ongoing B-cell aplasia in the spleen and BM of the mice, but reemergence of CD19+ cells in the CNS (**Fig. 2C-G**). This was seen in the presence of CAR T-cells, similar to the observation from our patient. We did not observe differences in the CD4/CD8 ratios and the memory profile of the CAR T-cells in the mice, though in all tissues there was a gradual expansion of CD4 cells compared to CD8 cells (**supplementary Fig. 3**).

**Figure 2:**
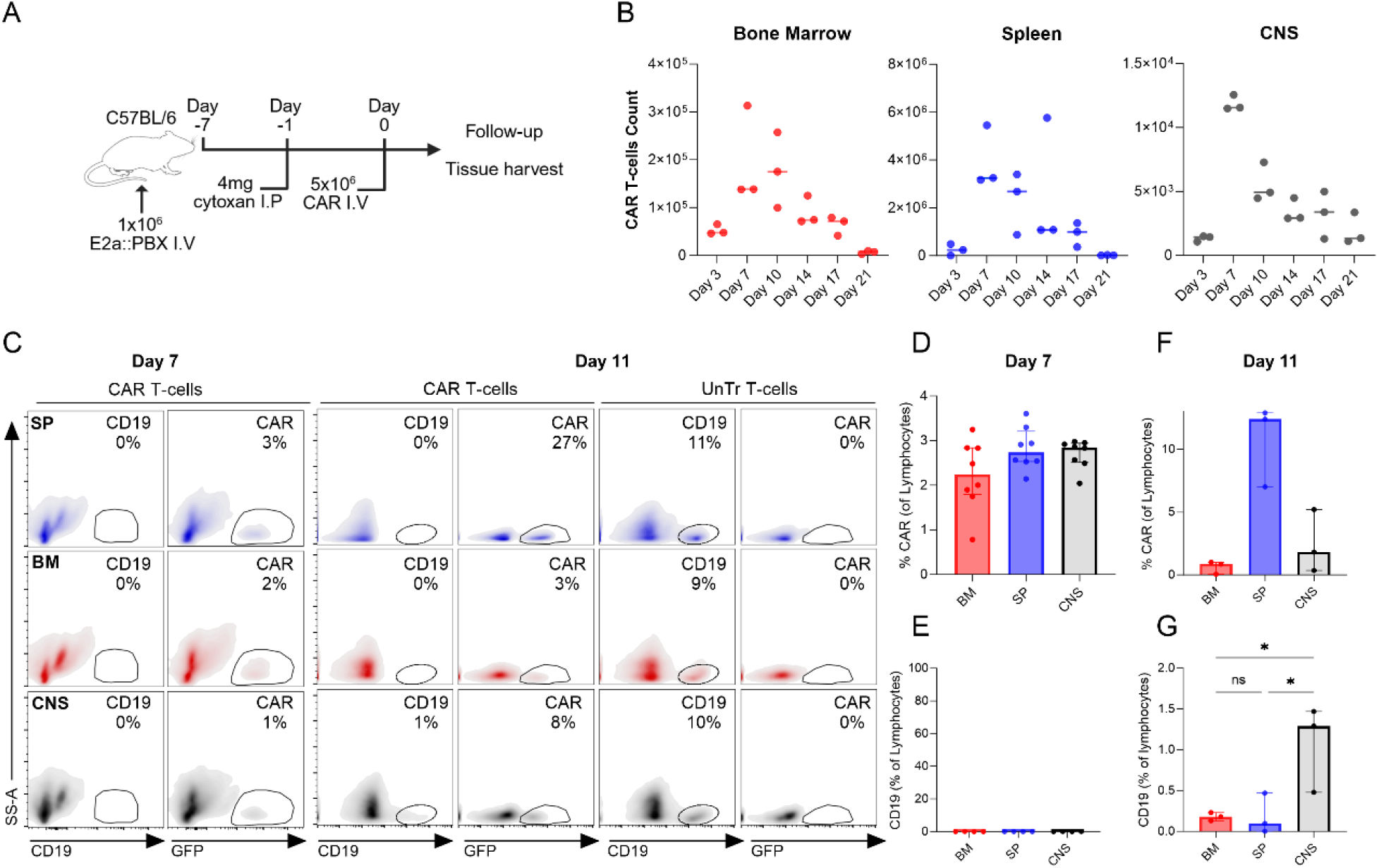
Murine model recapitulates CNS-specific CD19 positive emergence. **A** Schematic representation of the CD19 CAR-GFP murine experiment. C57BL/6 were treated with murine CD19 CAR-GFP 7 days after leukemia injection and 1 day after lymphodepletion with 4mg of cyclophosphamide. **B** mCD19 CAR-GFP live count as detected in flow cytometry, showing infiltration kinetics to bone marrow (red), spleen (blue) and central nervous system (CNS, black). **C-G** Enumeration of CAR+ and CD19+ cells in this model at two time points: **C** Representative flow cytometry plots of CAR+ and CD19+ cells in bone marrow, spleen and CNS at day 7 (left) and at day 11 (right) of C57Bl/6 mice treated according to the panel A with either CD19 CAR T-cells or untransduced T-cells (UnTr). Percentage of CAR T-cells (**D** and **F**) and CD19-cells (**E** and **G**) on day 7 and day 11 in the different tissues are shown. For all plots, error bars represent interquartile range. *P* values determined by ordinary One-Way ANOVA, \**p* ≤ 0.05.

We further assessed functionality of CAR T-cells harvested from different tissues in the mice. Day 7 MACS-sorted CAR^+^ cells from the spleen, BM and CNS were co-cultured with E2a::PBX1 leukemia at an effector to target (E:T) ratio of 1:8 for 48 hours (**Fig. 3a**). Leukemia killing was impaired when the CAR T-cells originated from the CNS, in comparison to CAR T-cells originating from the BM or spleen (**Fig. 3b-c)**. In addition, the number of CAR T-cells at the end of the culture was significantly lower when CAR T-cells from the CNS were used (**Fig. 3D-E)**. This was confirmed at an E:T ratio of 1:4 (**Supplementary Fig. 4)**. To further validate these findings, murine CAR T-cells were sorted based on GFP expression from all 3 tissues. Again, the CAR T-cells sorted from the CNS had inferior killing compared to those sorted from the spleen (**Supplementary Fig. 5)**.

**Figure 3:**
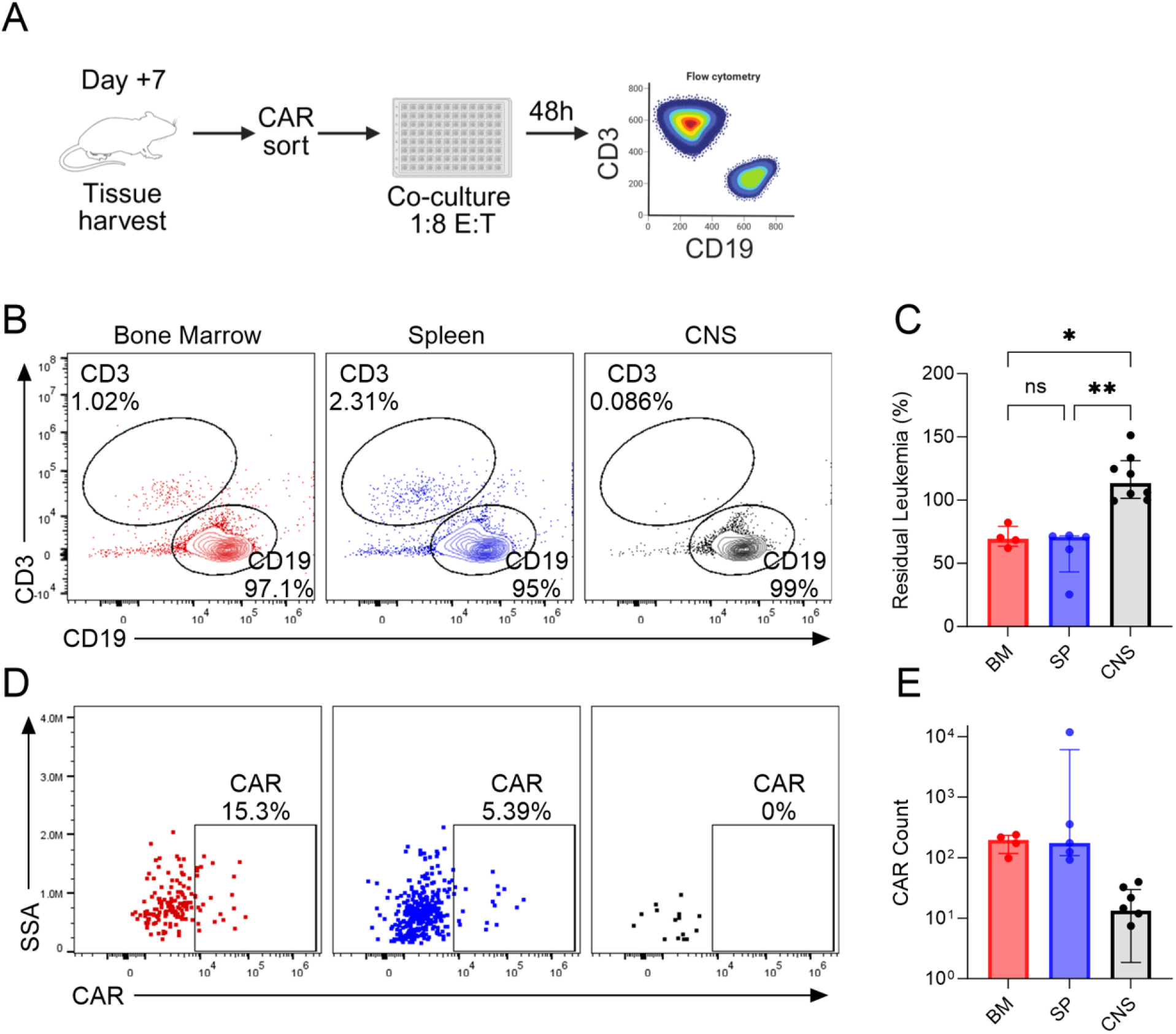
CNS-sorted murine CAR T-cells from are less cytotoxic compared to bone marrow and spleen CAR T-cells. **A** Schematic representation of the *ex vivo* killing experiment. C57Bl/6 mice were treated as in Fig. 2A. On day 7, cells from bone marrow (BM), spleen and central nervous system (CNS) were MACS-sorted for CAR+ expression, and cultured at a 1:8 effector to target (E:T) ratio with E2a::PBX CD19+ cells for 48 hours, then stained with CD19 and CD3 for flow cytometry and counted. **B** Representative flow cytometry plots of CD3+ and CD19+ cells after 48 hours of co-culturing E2a::PBX cells with CAR-T cells from BM (red), spleen (SP, blue) and CNS (black). **C** Cytotoxicity of sorted CAR-T cells was measured by number of residual CD19+ leukemic cells at the end of co-culture, normalized to the number of target cultured with no effector. **D** Representative flow cytometry plots of CAR-T cells gated on CD3+ cells from cells from the BM (red), spleen (blue) and CNS (black), after 48h co-culture with CD19+ target. **E** CAR-T cells live count at the end of the co-culture. For all plots, error bars represent interquartile range. *P* values determined by Kruskal-Wallis test, ns=non-significant (p > 0.05), \**p* ≤ 0.05, \*\**p* ≤ 0.01.

To confirm our findings in human CAR T-cells, we used NSG mice and injected them with 1×10^6^ Nalm6 leukemia cells, followed by human CAR T cells IV injection, without conditioning^17^. CAR T-cell expansion was evident in the BM, spleen and CNS of the mice, along with clearance of Nalm6 cells (**Supplementary Fig. 6**). Functional assessment of CAR T-cells from each tissue was carried out after MACS sorting for human CD8+, followed by a 48 hour co-culture at a 1:2 E:T ratio. Human CAR T-cells sorted from the CNS of the mice again demonstrated inferior killing poor expansion ex-vivo, compared to T-cells from the spleen or BM (**Fig. 4**). Altogether, this confirmed that CNS-residing murine and human CAR T-cells have impaired functionality *in vivo* and *ex-vivo*, compared to CAR T-cells harvested from the periphery.

**Figure 4:**
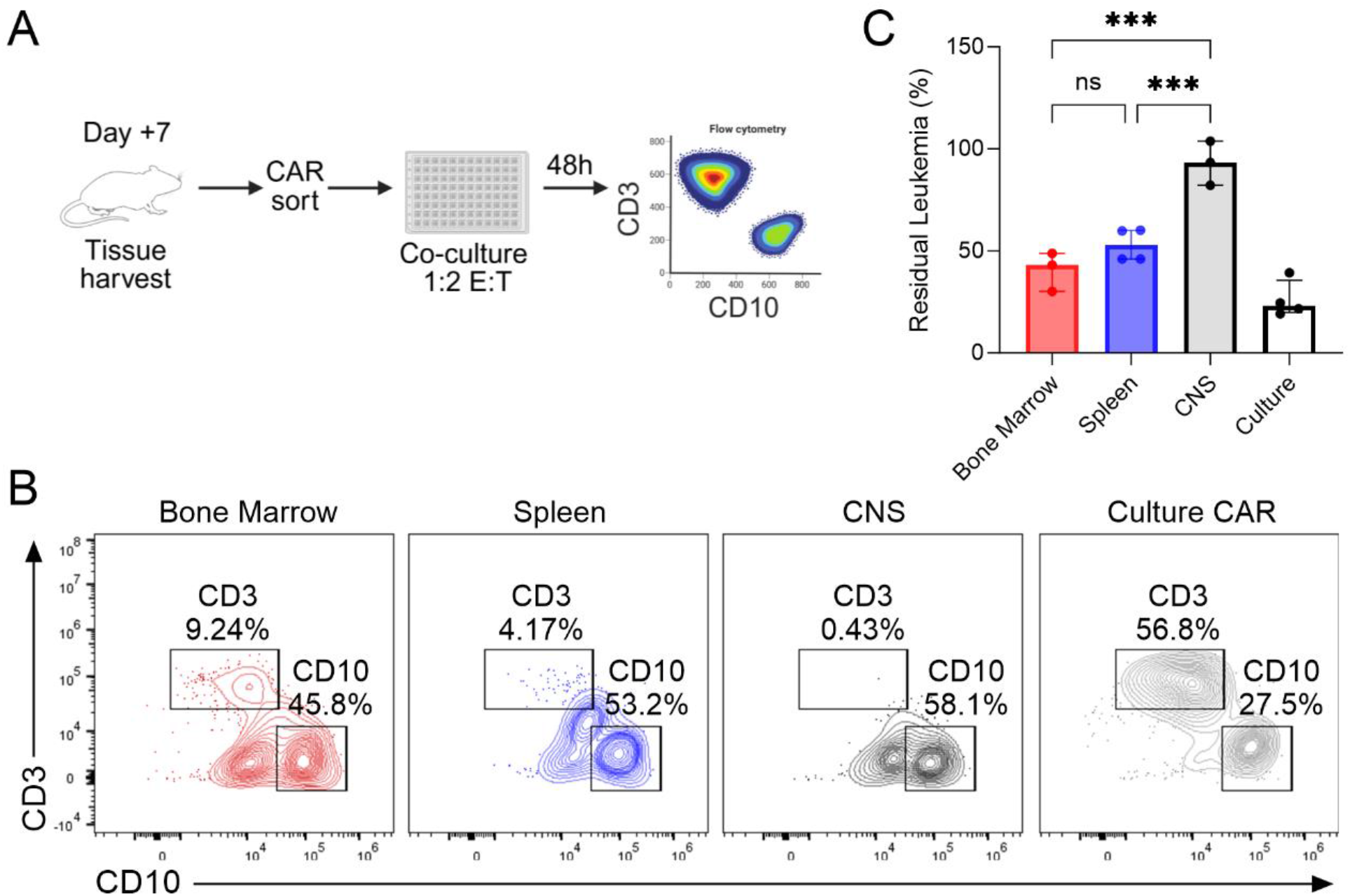
CNS-sorted huCAR from NSG mice are less cytotoxic *ex vivo* compared to sorted BM and Spleen CAR T-cells. **A** Schematic representation of the *ex vivo* killing experiment: NSG mice were injected with human CD19-28z CAR T-cells 1 week after leukemia inoculation. After 7 days, cells were harvested from the bone marrow, spleen and central nervous system (CNS) as well as CAR T-cells from original culture were MACS-sorted for human CD8+ and *ex vivo* cultured at a 1:2 effector to target ratio (E:T) with Nalm6 CD19+ cells for 48 hours, then stained with CD10 and CD3. **B** Representative flow cytometry plots of CD3+ and CD10+ cells after 48h co-culture of Nalm6 with CAR-T cells from the bone marrow (red), spleen (blue), CNS (black) and original culture (grey). **C** Cytotoxicity of sorted CAR was measured by number of residual leukemic cells at the end of co-culture, normalized to the number of target cells cultured with no effector. For all plots, error bars represent interquartile range. *P* values determined by ordinary One-Way ANOVA test, ns=non-significant (p > 0.05), \*\*\**p* ≤ 0.005.

One potential explanation for the poor functionality of CNS-residing CAR T-cells is their environment. The CSF is known to be poor in nutrients and oxygen compared to other tissues^21^. To study functionality of CAR T-cells in the CSF, human CAR T-cells were cultured in CSF, serum and culture media. The CSF and serum were paired, and originated from patients who were assessed routinely >28 days after CAR-T cell infusion on our clinical trial^22^, and >14 days after resolution of all CAR associated toxicities. CAR T-cells were thawed and recovered in medium containing 100 IU/ml IL-2 for 48 hours, and then transferred to unsupplemented CSF, serum or culture medium. In all conditions, CAR T-cells continued to expand, but expansion was slower in the CSF (**Fig. 5a-b**). Viability was similar between CSF and serum (**Fig. 5c)**, suggesting that the CSF does not lead to cell death but limits the proliferation of the cells. To test functionality in these media, CAR T-cells were co-cultured with Nalm6 cells at an 1:2 E:T ratio for 24 hours. Killing was assessed by enumerating residual Nalm6 cells compared to survival of the target cells in the same media in absence of CAR T-cells. When co-cultured in CSF, CAR T-cells had inferior killing compared to serum and culture media (**Fig. 5d-e)**.

**Figure 5:**
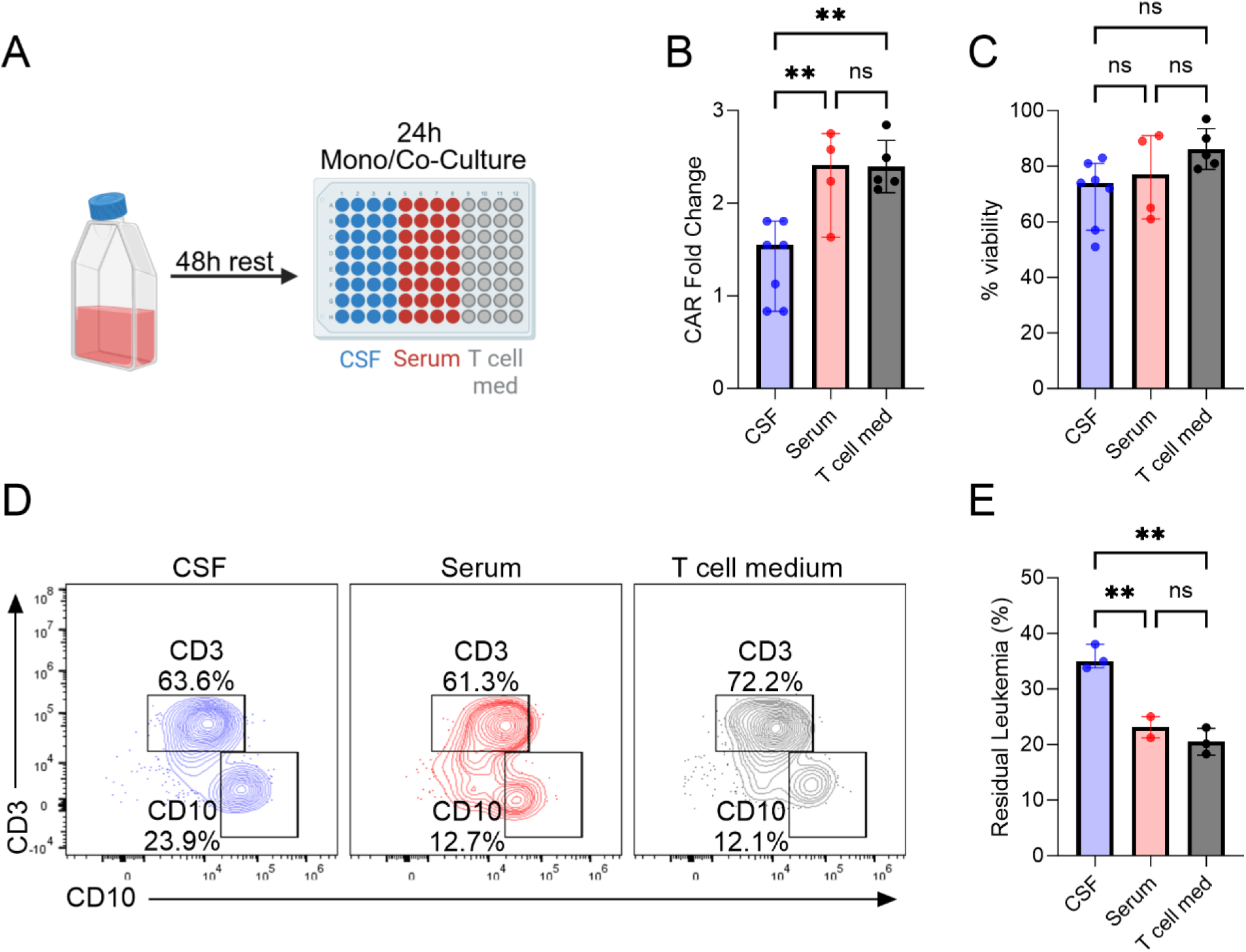
CD19-28z proliferation and cytotoxicity are impaired in human CSF compared to human Serum and T cell medium. **A** Schematic representation of the experiment: Human CD19-28z CAR T-cells were thawed and recovered in IL-2 supplemented T-cell medium for 48 hours prior to further experiments in un-supplemented human cerebral spinal fluid (CSF), matched human serum as well as T-cell medium. **B-C** CAR T-cells fold change growth (**B**) and viability (**C**) after 24h culture in 2 different matched human CSF and Serum and T-cell medium. **D-E** Recovered CAR-T cells were co-cultured with Nalm6 cells for 24 hours at a 1:2 effector to target ratio in either human CSF, matched serum, or T-cell media. D Representative flow cytometry plots of CD3+ and CD10+ cells at the end of the co-culture. **E** Cytotoxicity in different human mediums was measured by number of residual leukemic cells, normalized to target with no effector grown in each of the media. For all plots, error bars represent interquartile range. *P* values determined by ordinary Kruskal-Wallis test, ns=non-significant (p > 0.05), \*\**p* ≤ 0.01.

Another possible corroborator to the impaired functionality of the CAR T-cells in the CNS is the BBB, designed to protect the brain from intravascular substances and toxins^23^. To better characterize BBB-migrating CAR T-cells and the cross talk of migrating T-cells with the BBB, we utilized a unique model of human BBB^19,23^, consisting of a transwell layered with CD34+ human endothelium and bovine pericytes, leading to “blood” (luminal) and “brain” (abluminal) compartments (**Fig. 6a**). We confirmed CAR T-cells ability to migrate across the BBB-model. The migration was unrelated to the presence of target in the abluminal compartment (**Supplementary Fig. 7a**) but was substantially reduced when the CAR T-cells were preactivated with Nalm6 cells (**Supplementary Fig. 7b**). After allowing for a 4-hour migration time the number of CAR+ cells was similar in both compartments, although there was a slight decrease in the percentage of CAR+ cells in the abluminal compartment (**Fig. 6B-C**). CAR T-cells from the abluminal (“brain”) side were activated in the absence of target, showing higher CD69 expression, but lower or almost no 4-1BB expression (**Fig. 6D-E**). CAR T-cells were collected from the luminal and abluminal sides of the BBB-model, and co-cultured at a 1:2 E:T ratio for 24 hours. The abluminal CAR T-cells were less cytotoxic compared to CAR T-cells from the luminal side and CAR T-cells from the original culture (**Fig. 6F-G**), though IFNγ production was increased in the abluminal CAR T-cells compared to the luminal CAR T-cells (**Fig. 6H)**. Of note, exhaustion markers (PD-1, TIM3 and LAG3) were not increased in CAR T-cells at the abluminal compartment compared to the luminal side, though both were higher than culture CAR T-cells (**Supplementary Fig. 8**). The increase of CD69 and IFNγ production in the face of low 4-1BB and reduced killing suggested that the BBB-crossing CAR T-cells underwent activation induced cell death (AICD). Indeed, Annexin binding was significantly higher in BBB-crossing CAR T-cells compared to the luminal CAR T-cells and those grown in culture (**Fig. 6I-K**).

**Figure 6:**
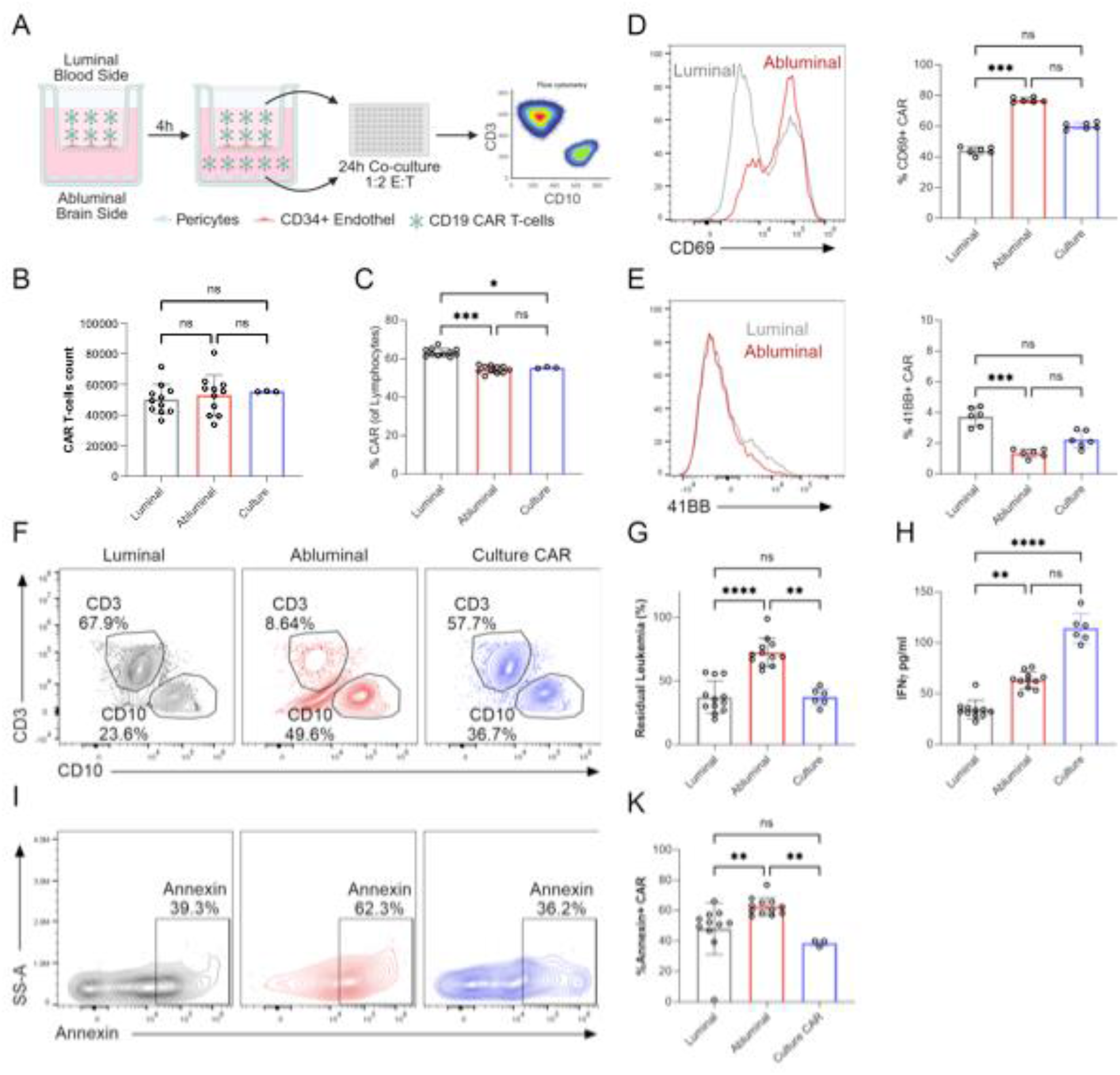
BBB-migrated CD19 CAR T-cells are less cytotoxic against leukemic cells and undergo AICD. **A** Schematic representation of *in vitro* human BBB model migration experiment: Human CD19 CAR T-cells were introduced to the luminal (blood) side of a transwell (TW) insert mimicking human BBB composed of human CD34+ endothelial cells and bovine brain pericytes. Following 4 hours of migration, cells were collected from both luminal and abluminal (brain) compartments and were either analyzed for phenotype or co-cultured with Nalm6 CD19+ cells at a 1:2 effector to target ratio (E:T) for 24 hours. **B-C** CAR T-cells live count (**B**) and percentage of CAR+ cells of total cells collected (**C**) from luminal, abluminal and original culture. **D** Representative histogram (left) and percentage (right) of CD69 expression in CAR T-cells collected from luminal and abluminal. **E** Representative histogram (left) and percentage (right) of 41BB expression in CAR T-cells collected from luminal and abluminal. **F** Representative flow cytometer plot of CD3+ and CD10+ cells after 24h co-culture of the collected CAR T-cells with Nalm6. **G** Cytotoxicity of CAR T-cells from different compartments was measured by number of residual leukemic cells, normalized to the number of target cells grown with no effector. **H** Levels of IFNγ in the supernatant collected at the end of a 24h co-culture. **I** Representative flow cytometry plots of Annexin^+^ binding gated on CAR^+^ T-cells from luminal, abluminal and culture. **K** Percentage of Annexin^+^-binding CAR T-cells. For all plots, error bars represent interquartile range. *P* values determined by ordinary Kruskal-Wallis test, ns=non-significant (p > 0.05), \**p* ≤ 0.05, \*\**p* ≤ 0.01, \*\*\**p* ≤ 0.005.

Experiments were carried out with CD28-costimulated CAR T-cells, associated with increased AICD^24^. We confirmed our findings using 4-1BB costimulated CD22 CAR T cells, to control both for different binding and different costimulation. CD22 CAR T cells that migrated across the BBB model also showed increased annexin binding compared to the luminal and culture controls and subsequent poor killing of leukemia (**Fig. 7**).

**Figure 7:**
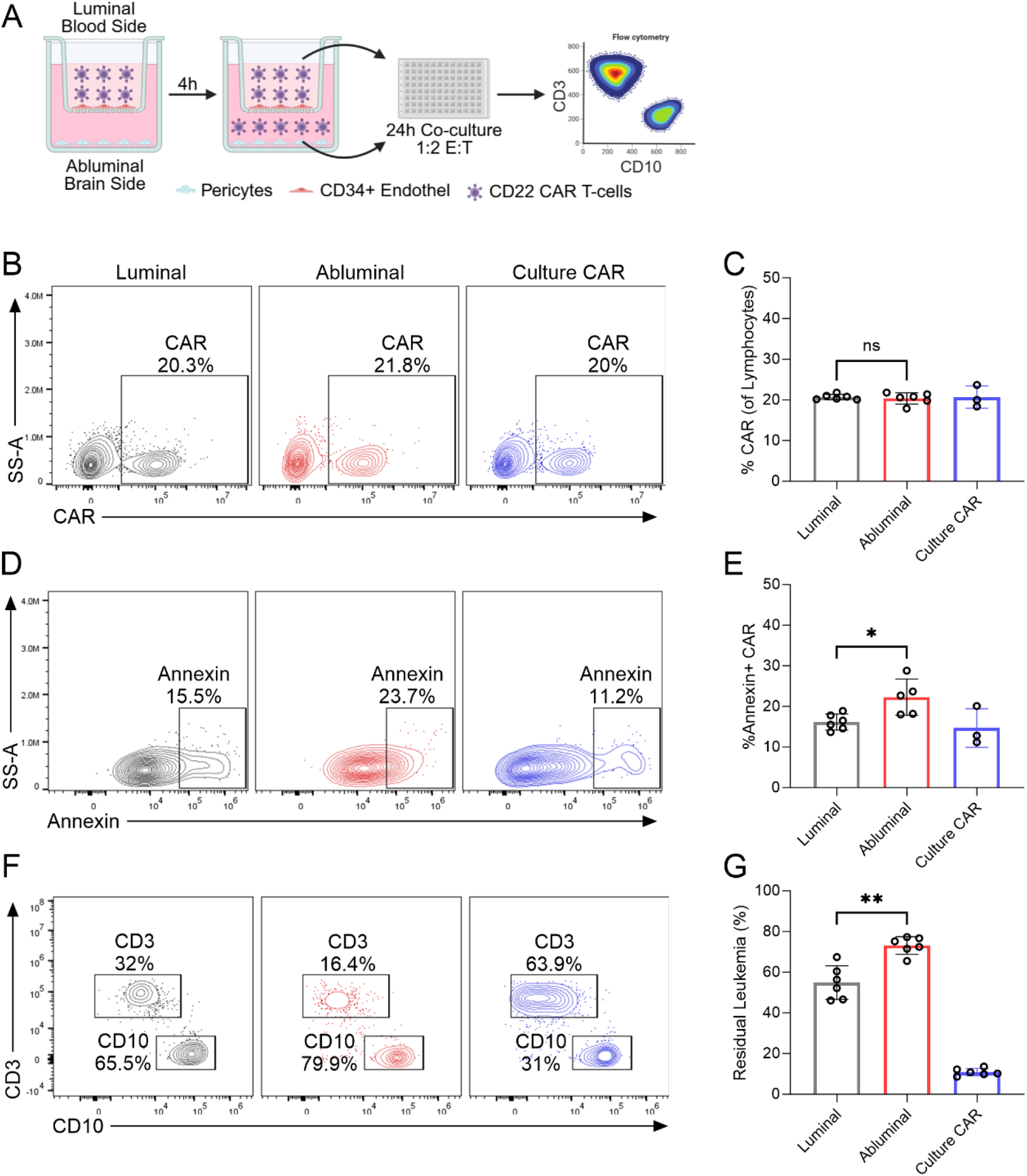
BBB-migrated CD22 CAR T-cells are less cytotoxic against leukemic cells and undergo AICD. **A** Schematic representation of *in vitro* BBB model migration experiment: Human CD22 CAR T–cells (m971-41BB-CD3z) were introduced to the luminal (blood) side of a transwell (TW) insert mimicking human BBB (as in Fig.6), and allowed to migrate for 4 hours. Cells were collected from both luminal and abluminal (brain) sides and either analyzed by flow cytometry or co-cultured with Nalm6 CD22+ cells at a 1:2 effector to target ratio (E:T) for 24 hours. **B** Representative flow cytometry plots of CAR T-cells from luminal, abluminal and culture. **C** CAR percentage from total cells collected from luminal, abluminal and original culture. **D** Representative flow cytometry plots of Annexin^+^ binding gated on CAR+ T-cells from luminal, abluminal and culture. **E** percentage of Annexin^+^-binding CAR T-cells. **F** Representative flow cytometry plots of CD3+ and CD10+ cells after 24h co-culture of Nalm6 with CAR T-cells from all compartments. **G** Cytotoxicity of CAR T-cells from different compartments was measured by number of residual leukemic cells, normalized to the number of target cells grown with no effector. For all plots, error bars represent interquartile range. *P* values determined by ordinary Kruskal-Wallis test, ^ns^*p* > 0.05, \**p* ≤ 0.05, \*\**p* ≤ 0.01.

Previous studies suggested CD19 expression on pericytes as a potential cause for CAR-associated neurotoxicity^25^. To confirm that activation and AICD is not related to expression on subsets of the model, after assembly of the BBB, we cultured CAR T-cells separately with CD34+ human endothelium and bovine pericytes, in addition to the controls in the luminal and abluminal compartments (**supplementary Fig.9A-B**). We confirmed AICD by annexin binding **supplementary Fig.9C-D**) and poor activity (**supplementary Fig. 9E-F**) occur in the abluminal migrating CAR T-cells and not with other conditions.

## Discussion

With the increased clinical use of CAR T-cells in hematological malignancies and beyond, concerns regarding resistance mechanisms are emerging. While antigen-positive relapse is usually associated with loss of the CAR T-cell persistence^26^, the presence of CD19^+^ relapse confined to the CNS in patient treated with CD19 CAR T-cells, while in the peripheral blood there is evidence of ongoing CAR T-cell activity, suggests other mechanisms. This eludes that the CNS is a hostile environment to CAR T-cells. Indeed, there are reports of success of CAR T-cells in eradicating disease in the CNS but concerns regarding durability of these remissions are arising^8,10^.

In this work we generated mouse models and confirmed inferior efficacy of CAR T-cells derived from the CNS compared to other hematopoietic tissues, in both immunocompetent and humanized murine models. We sought to identify the mechanisms contributing to this dysfunction and identified two-the CSF environment and the BBB.

Initially, we examined the effect of the CSF on CAR T-cells function. A recent report focused on the impact of the CSF on CAR T-cells^27^. Overall, the CSF is low in nutrients and may thus impair immune cell function. The different metabolic composition of the CSF compared to the peripheral blood (PB), including increased TCA-cycle metabolites and reduced lipids, impacted the immunometabolic features of CAR T-cells, increasing LAL-mediated lipolysis^27^. In our experiments, using patient-derived CSF and paired serum controls, CAR T-cells had impaired growth in CSF but preserved the ability to kill target cells. However, the killing was significantly inferior when compared to other mediums. Other work suggested that culture in the CSF may induce improved properties to CAR T-cell function^27^. We did not attempt prolonged culture of the cells in CSF, but our data did show limited growth in the CSF without decreased viability. Thus, the improved outcome reported in CSF-cultured CAR T-cells^27^ may be a result of halting initial expansion, as seen in other models with signaling blockade^28^ or metabolic manipulation^29^ showing improved functionality of these less-expanded cells.

Our work specifically examined the contribution of migration through the BBB on CAR T-cell function. Using a human *in vitro* BBB model, we found that CAR T-cells that migrated across the BBB showed reduced cytotoxicity compared to non-migrating or resting cells. We identified that CAR T-cells that crossed the BBB model exhibited a unique activation profile despite the absence of target leukemic cells, including increased CD69 expression, increased IFNγ production, and similar low 4-1BB expression. Other work has reported increased CD69 expression on CAR T-cells from the CSF compared to the peripheral blood on day +28 after infusion^27^, while CD19+ cells were absent in all compartments, further confirming antigen-independent activation in this compartment. Our findings that the BBB-migrating CAR T-cells are less cytotoxic, express activation markers and our preliminary findings that activated CAR T-cells were less abundant upon crossing the BBB model – were concerning for AICD. Indeed, migrating CAR T-cells had an increased apoptotic state compared to non-migrating and resting CAR T-cells. 4-1BB costimulation is associated with reduced AICD^30^, and improved survival of CAR-T cells^24^. We confirmed our findings to be consistent across both CD28- and 4-1BB-costimulated CAR constructs, and reproduced with CD22 targeting, indicating that the dysfunction of BBB-crossing cells is not antigen-specific nor limited to a particular co-stimulatory domain. Importantly, we ruled out non-specific activation due to antigen expression on BBB components, as co-culture with either CD34+ endothelial cells or pericytes alone did not reproduce the same phenotype, supporting the notion that crossing the full BBB apparatus is required to trigger activation and further AICD in CAR T-cells.

This work focused on CAR T-cells, but may be relevant to other T-cell engaging therapies. A recent study in children with ALL demonstrated that the addition of blinatumomab, a bispecific CD19-CD3 T-cell engager, to consolidation therapy in ALL significantly reduced relapse risk, but did not impact the risk of CNS relapse^31^. We have previously shown that T-cells are more abundant in the CNS after blinatumomab or CAR T-cells^32^. Thus, there may be a functional deficit in clearing leukemia with immune targeting in the CNS. The phenotype and activity of T-cells from patients treated with these therapies should be the subject of future studies.

Our study has several limitations. While our murine and humanized models allow functional interrogation of CAR T-cells derived from the CNS, it lacks the ability to interrogate the fully complex human CNS immune microenvironment and the role of other BBB and brain resident cells such as microglia and astrocytes. Second, our *in vitro* BBB model does not account for dynamic physiological factors such as shear stress from blood flow or long-term interactions with CNS-resident immune and stromal cells. Future experiments should also investigate other downstream apoptotic pathways associated with BBB-induced AICD, as well as BBB-specific properties influencing CAR T-cells activation and survival.

In summary, we show data from *in vivo* models, *in vitro* models, and a patient case indicating relative dysfunction of CAR T-cells in the CNS, and suggest two corroborating mechanisms: the nutrient-poor environment in the CSF, and a unique BBB-associated dysfunction. With the increase use of CAR T-cells for disorders of the CNS, direct injection may overcome some of these obstacles, and is currently tested in brain tumors and in CNS lymphoproliferative disorders. Further work focusing on studying CNS-residing T-cells from patients, as well as minimizing AICD (e.g. through IFNγ inhibition), will be essential to validate the clinical relevance of these mechanisms.

## Acknowledgements

The authors would like to thank James Kochenderfer, Terry Fry and Naomi Taylor (NCI) for providing the murine CAR construct and the murine leukemia. We would like to thank Jonathan Esenstein and Gal Cafri for their assistance with the CD22 CAR, and Tal Yardeni and Shani K Liskov for assistance and advice in mouse experiments. This work was funded by the Dotan center for hematologic malignancies grant (EJ and IC) and the Israel Cancer Association Grant 20211196 (EJ). This work was carried out in partial requirements for YA PhD degree (Gray Faculty of Medical and Health Sciences, Tel Aviv University).

## Author Contribution

YAT, SI and EJ conceived the study plan. YAT, OR, AM, HL, DB and AA carried out in vitro and in vivo experiments. YAT, OR and IC designed and carried out the in vitro BBB model experiments. ADY, TF and SI provided clinical data on the patient presented. YAT and EJ wrote the first version of the manuscript. All authors edited and approved the final version.

## Data Availability Statement

All data generated or analyzed for this study are included in this article (and its supplementary information files).

## References

1. Thastrup M, Duguid A, Mirian C, Schmiegelow K, Halsey C. Central nervous system involvement in childhood acute lymphoblastic leukemia: challenges and solutions. Leukemia 2022; : 1–18.

2. Pui C-H, Yang JJ, Hunger SP, Pieters R, Schrappe M, Biondi a. et al. Childhood Acute Lymphoblastic Leukemia: Progress Through Collaboration. J Clin Oncol 2015; 33. doi:10.1200/JCO.2014.59.1636.

3. Frishman-Levy L, Izraeli S. Advances in understanding the pathogenesis of CNS acute lymphoblastic leukaemia and potential for therapy. Br J Haematol 2017; 176: 157–167.

4. Rheingold SR, Bhojwani D, Ji L, Xu X, Devidas M, Kairalla JA et al. Determinants of survival after first relapse of acute lymphoblastic leukemia: a Children’s Oncology Group study. Leukemia 2024. doi:10.1038/s41375-024-02395-4.

5. Houillier C, Soussain C, Ghesquières H, Soubeyran P, Chinot O, Taillandier L et al. Management and outcome of primary CNS lymphoma in the modern era. Neurology 2020; 94. doi:10.1212/WNL.0000000000008900.

6. Jacoby E, Shahani SA, Shah NN. Updates on CAR T-cell therapy in B-cell malignancies. Immunol Rev 2019; 290: 39–59.

7. Frigault MJ, Dietrich J, Martinez-Lage M, Leick M, Choi BD, DeFilipp Z et al. Tisagenlecleucel CAR-T Cell Therapy in Secondary CNS Lymphoma. Blood 2019;: blood.2019001694.

8. Jacoby E, Ghorashian S, Vormoor B, De Moerloose B, Bodmer N, Molostova O et al. CD19 CAR T-cells for pediatric relapsed acute lymphoblastic leukemia with active CNS involvement: a retrospective international study. Leukemia 2022; 36: 1525–1532.

9. Fabrizio VA, Phillips CL, Lane A, Baggott C, Prabhu S, Egeler E et al. Tisagenlecleucel outcomes in relapsed/refractory extramedullary ALL: A Pediatric Real World CAR Consortium Report. Blood Adv 2022; 6: 600–610.

10. Ossami Saidy A, Peczynski C, Thieblemont C, Daskalakis M, Wehrli M, Beauvais D et al. Efficacy and safety of CAR T-cell therapy in patients with primary or secondary CNS lymphoma: A study on behalf of the EBMT and the GoCART coalition. HemaSphere 2025; 9: e70146.

11. Epperla N, Feng L, Shah NN, Fitzgerald L, Shah H, Stephens DM et al. Outcomes of patients with secondary central nervous system lymphoma following CAR T-cell therapy: a multicenter cohort study. J Hematol Oncol 2023; 16: 111.

12. O’Rourke DM, Nasrallah MP, Desai A, Melenhorst JJ, Mansfield K, Morrissette JJD et al. A single dose of peripherally infused EGFRvIII-directed CAR T cells mediates antigen loss and induces adaptive resistance in patients with recurrent glioblastoma. Sci Transl Med 2017; 0984.

13. Choi BD, Gerstner ER, Frigault MJ, Leick MB, Mount CW, Balaj L et al. Intraventricular CARv3-TEAM-E T Cells in Recurrent Glioblastoma. N Engl J Med 2024; 390: 1290–1298.

14. Majzner RG, Ramakrishna S, Yeom KW, Patel S, Chinnasamy H, Schultz LM et al. GD2-CAR T cell therapy for H3K27M-mutated diffuse midline gliomas. Nature 2022; 603: 934–941.

15. Jacoby E, Chen A, Loeb DM, Gamper CJ, Zambidis E, Llosa NJ et al. Single-Agent Post-Transplantation Cyclophosphamide as Graft-versus-Host Disease Prophylaxis after Human Leukocyte Antigen-Matched Related Bone Marrow Transplantation for Pediatric and Young Adult Patients with Hematologic Malignancies. Biol Blood Marrow Transplant 2016; 22: 112–118.

16. Rozenbaum M, Fluss R, Marcu-Malina V, Sarouk I, Meir A, Elitzur S et al. Genotoxicity Associated with Retroviral CAR Transduction of ATM-Deficient T Cells. Blood cancer Discov 2024; 5: 267–275.

17. Rozenbaum M, Meir A, Aharony Y, Itzhaki O, Schachter J, Bank I et al. Gamma-Delta CAR-T Cells Show CAR-Directed and Independent Activity Against Leukemia. Front Immunol 2020; 11: 1–8.

18. Jacoby E, Yang Y, Qin H, Chien CD, Kochenderfer JN, Fry TJ. Murine allogeneic CD19 CAR T-cells harbor potent anti-leukemic activity but have the potential to mediate lethal GVHD. Blood 2016; 127: 1361–1370.

19. Cecchelli R, Aday S, Sevin E, Almeida C, Culot M, Dehouck L et al. A stable and reproducible human blood-brain barrier model derived from hematopoietic stem cells. PLoS One 2014; 9: e99733.

20. Shelly S, Liraz Zaltsman S, Ben-Gal O, Dayan A, Ganmore I, Shemesh C et al. Potential neurotoxicity of titanium implants: Prospective, in-vivo and in-vitro study. Biomaterials 2021; 276: 121039.

21. Savino AM, Fernandes SI, Olivares O, Zemlyansky A, Cousins A, Markert EK et al. Metabolic adaptation of acute lymphoblastic leukemia to the central nervous system microenvironment depends on stearoyl-CoA desaturase. Nat Cancer 2020; 1: 998–1009.

22. Jacoby E, Bielorai B, Hutt D, Itzhaki O, Adam E, Bar D et al. Parameters of long-term response with CD28-based CD19 chimaeric antigen receptor-modified T cells in children and young adults with B-acute lymphoblastic leukaemia. Br J Haematol 2022; 197: 475–481.

23. Ravid O, Elhaik Goldman S, Macheto D, Bresler Y, De Oliveira RI, Liraz-Zaltsman S et al. Blood-brain barrier cellular responses toward organophosphates: Natural compensatory processes and exogenous interventions to rescue barrier properties. Front Cell Neurosci 2018; 12: 1–14.

24. Long AH, Haso WM, Shern JF, Wanhainen KM, Murgai M, Ingaramo M et al. 4-1BB costimulation ameliorates T cell exhaustion induced by tonic signaling of chimeric antigen receptors. Nat Med 2015; 21: 581–590.

25. Parker KR, Migliorini D, Perkey E, Yost KE, Bhaduri A, Bagga P et al. Single-Cell Analyses Identify Brain Mural Cells Expressing CD19 as Potential Off-Tumor Targets for CAR-T Immunotherapies. Cell 2020; 0: 1–17.

26. Shah NN, Fry TJ. Mechanisms of resistance to CAR T cell therapy. Nat. Rev. Clin. Oncol. 2019; 16: 372–385.

27. Goldberg L, Haas ER, Urak R, Vyas V, Pathak K V., Garcia-Mansfield K et al. Immunometabolic adaptation of CD19-targeted CAR T cells in the central nervous system microenvironment of patients promotes memory development. Cancer Res 2024. doi:10.1158/0008-5472.CAN-23-2299.

28. Weber EW, Parker KR, Sotillo E, Lynn RC, Anbunathan H, Lattin J et al. Transient rest restores functionality in exhausted CAR-T cells through epigenetic remodeling. Science 2021; 372. doi:10.1126/science.aba1786.

29. Gross G, Alkadieri S, Meir A, Itzhaki O, Aharoni-Tevet Y, Ben Yosef S et al. Improved CAR-T cell activity associated with increased mitochondrial function primed by galactose. Leukemia 2024; 38: 1534–1540.

30. Stärck L, Scholz C, Dörken B, Daniel PT. Costimulation by CD137/4-1BB inhibits T cell apoptosis and induces Bcl-xL and c-FLIP(short) via phosphatidylinositol 3-kinase and AKT/protein kinase B. Eur J Immunol 2005; 35: 1257–66.

31. Gupta S, Rau RE, Kairalla JA, Rabin KR, Wang C, Angiolillo AL et al. Blinatumomab in Standard-Risk B-Cell Acute Lymphoblastic Leukemia in Children. N Engl J Med 2024; : 1–16.

32. Leib S, Bielorai B, Vernitsky H, Aharony-Tevet Y, Toren A, Jacoby E. Cerebral Spinal Fluid Parameters Following CD19-Targeted Therapies in Children and Young Adults. J Pediatr Hematol Oncol 2024; 46: 29– 32.

