## Supplemental Figures for "CAR T-cells dysfunction in the central nervous system is mediated by BBB-induced activation-induced cell death"

**CAR T-cells dysfunction in the CNS is mediated by BBB-induced AICD**

**Supplemental material**

**Table of contents**

**Table of flow antibodies**

anti whitlow/218 linker and anti G4S linker (Cell Signaling), CD19 CAR reagent and CD22 CAR reagent, anti-mouse CD19, anti-mouse CD3, anti-human CD10, anti-human CD3, anti-human LAG3, anti-human 41BB, Annexin-V (Miltenyi), anti-mouse CD4, anti-mouse CD8a, anti-mouse CD62L, anti-mouse CD44, anti-human TIM3, anti-human PD1, anti-human CD69, streptavidin (biolegend)

Supplementary Figure 1

Murine CD19-CAR GFP expression

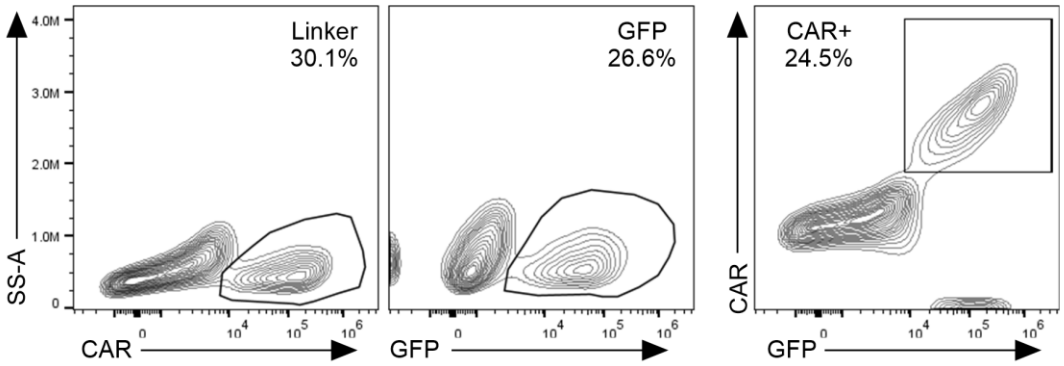


Flow cytometry representative plots of murine CAR T-cell expression measured by whitlow linker (left), GFP-expression (middle) and their correlation (right)

Supplementary Figure 2

Efficacy of murine CD19-CAR GFP$^+$ T-cells

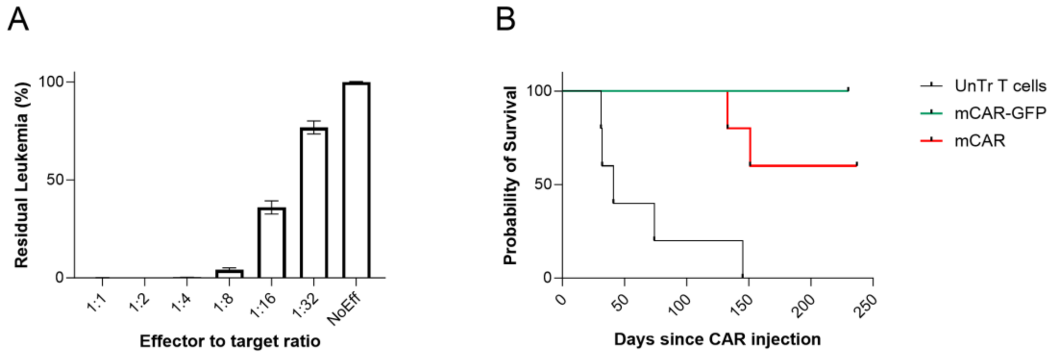


**A** Murine CD19-CAR GFP$^+$ T-cells were co-cultured with E2a::PBX CD19+ murine leukemic cells, at several effector to target ratios for 48 hours. Cytotoxicity was measured by number of residual leukemic cells at the end of co-culture, normalized to the number of target grown with no effector. **B** C57BL/6 were intravenously injected with E2a::PBX leukemia on day -7, cyclophosphamide (4mg/mouse) on day -1, and T-cells on day 0. Probability of survival mice treated with murine CD19 CAR T-cells (mCAR, red), CD19 CAR-GFP T-cells (mCAR-GFP, green) and untransduced T-cells (UnTr T cells, grey) is shown.

Supplementary Figure 3

Phenotype of murine CD19 CAR T cells harvested from different tissues

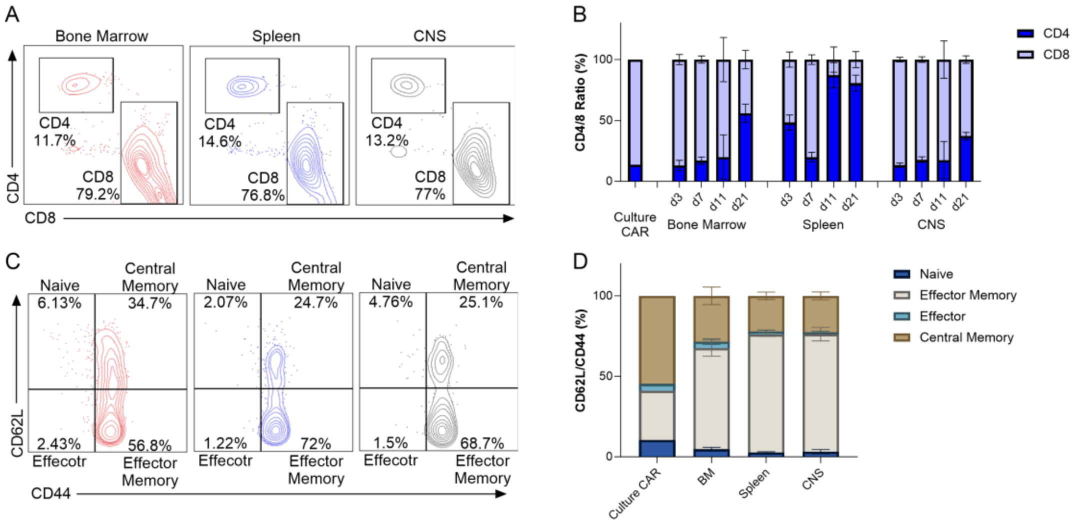


**A** Flow cytometry representative plots of CD4+ and CD8+ CAR T-cells from Bone Marrow, spleen and CNS of mCD19 CAR-GFP treated mice (as in Fig. 2A). **B** Percentage of CD4+ and CD8+ CAR T-cells from receptive tissues. **C** Flow cytometry representative plots of memory profile of CAR T-cells harvested on day 7 from murine tissues, based on CD62L and CD44 expression. **D** Percentage of memory profile of CAR T-cells from mice tissues.

Supplementary Figure 4

Cytotoxicity of MACS-sorted murine CAR T-cells from the CNS, BM and Spleen

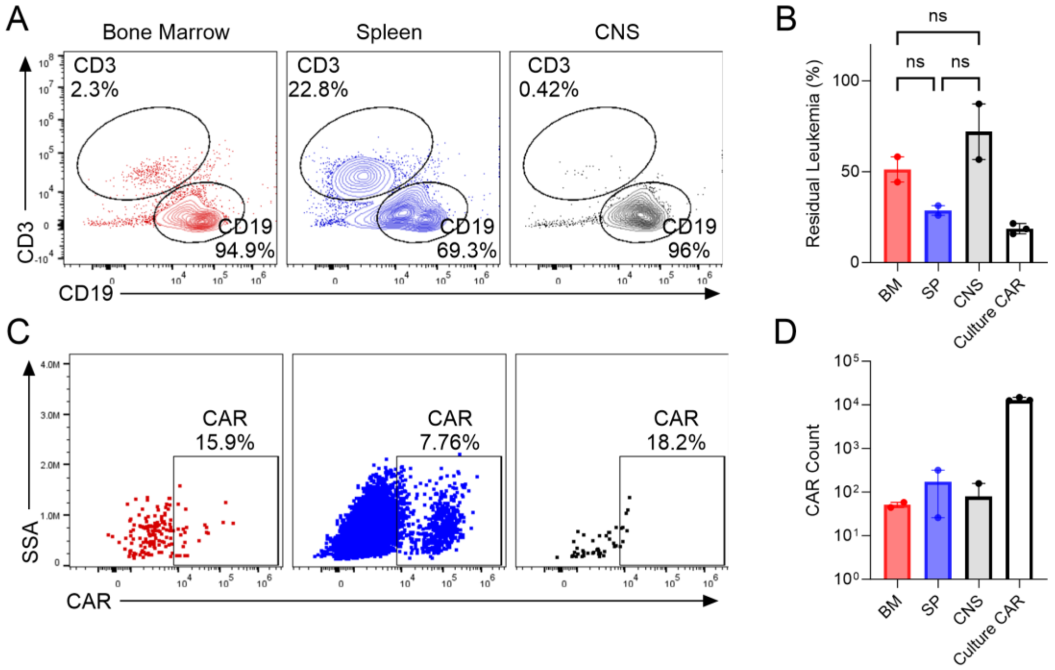


C57Bl/6 mice were treated as in Fig. 2A. On day 7, cells from bone marrow (BM), spleen and central nervous system (CNS) were MACS-sorted for CAR+ expression, and cultured at a 1:4 effector to target (E:T) ratio with E2a::PBX CD19+ cells for 48 hours. **A** representative flow cytometry plots of CD3+ and CD19+ cells after 48h co-culture. **B** cytotoxicity of sorted CAR was measured by percentage of residual CD19-leukemic cells at the end of co-culture, normalized to target with no effector. **C** representative flow cytometry plots of CAR T-cells gated on CD3+ cells showing proliferation of CAR T-cells from bone marrow and spleen and but not from CNS after 48h co-culture with CD19+ target. **D** CAR live count at the end of the co-culture. For all plots, error bars represent interquartile range. *P* values determined by Kruskal-Wallis test, $^{ns}p > 0.05$.

Supplementary Figure 5

Cytotoxicity of GFP-sorted CAR T-cells from the CNS, BM and Spleen

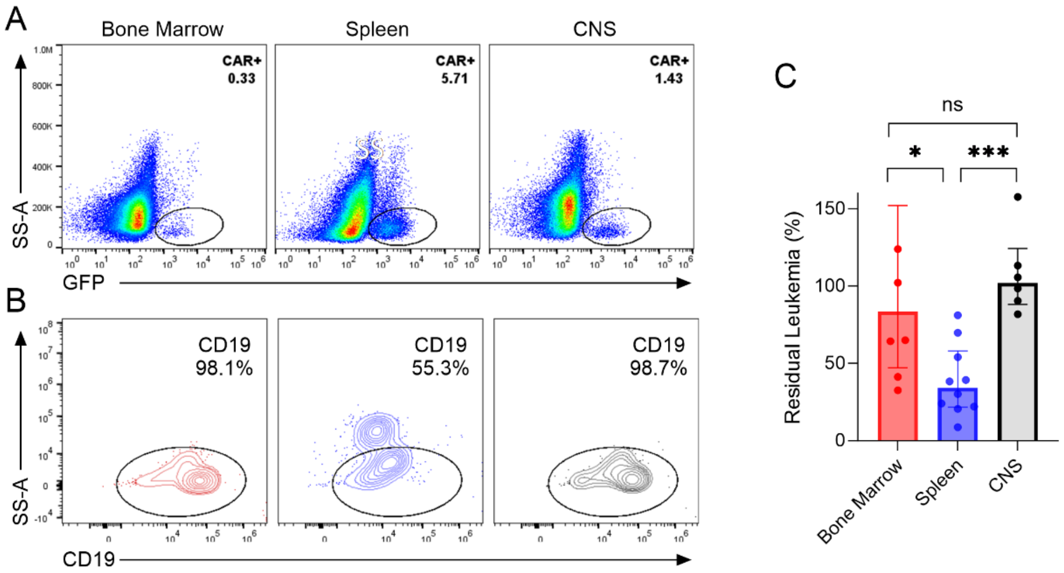


Experiment design was as in Fig. 2A and 3A. Here, CAR T-cells from tissues was sorted based on GFP expression of CAR+ cells. **A** Flow cytometry plots showing sort gates for murine CD19 CAR-GFP T-cells from bone marrow, spleen and CNS of mice. **B** Representative flow cytometry of CD3+ and CD19+ cells after 48h co-culture iat a 1:2 effector to target ratio. **C** Cytotoxicity of sorted CAR was measured by number of residual CD19-leukemic cells at the end of co-culture, normalized to target cultured with no effector. For all plots, error bars represent interquartile range. *P* values determined by Mann-Whitney test, $^{ns}p > 0.05$, $^{*}p \leq 0.05$, $^{***}p \leq 0.005$.

Supplementary Figure 6

Human CAR-T cell expansion in NSG mice

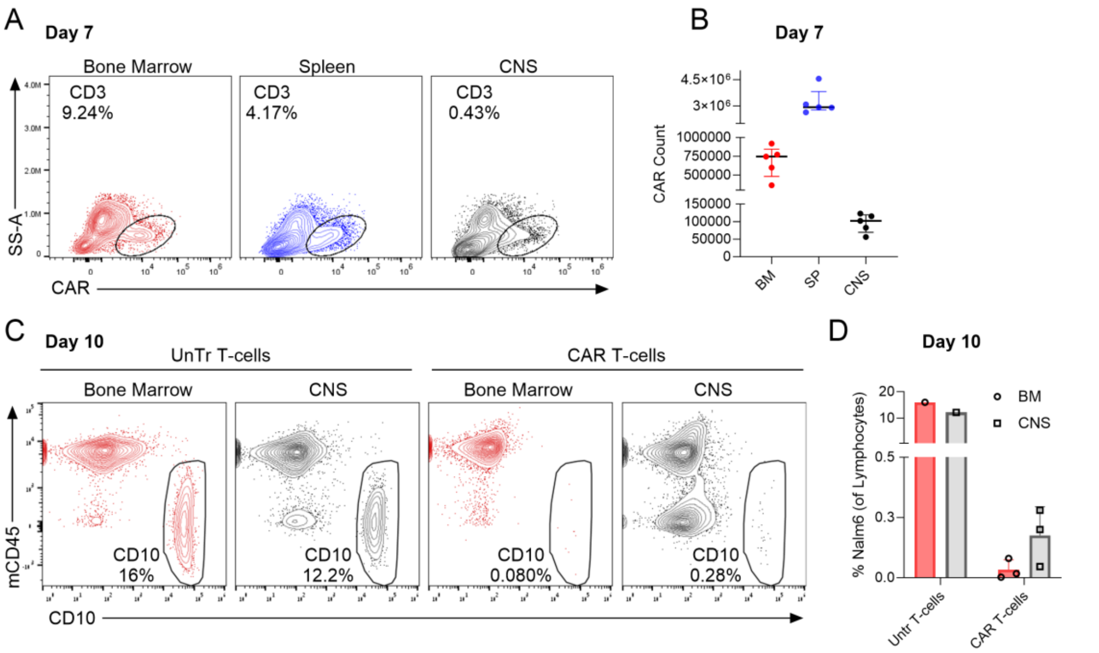


NSG mice were intravenously administered Nalm6 cells on day -7 and CD19 CAR or untransduced T-cells on day 0. **A** Flow cytometry plots showing infiltration of human CD19 CAR T-cells to bone marrow, spleen and CNS on day 7 post CAR T-cells injection. **B** CAR T-cell live count in bone marrow, spleen and CNS on day 7. **C** Flow cytometry plots of the bone marrow and CNS of the NSG mice on day +10 post untransduced T-cells (left) CAR T-cells (right) injection. **D** Percentage of Nalm6 cells in bone marrow (BM) and CNS of untrasnduced and CAR T-cells treated NSG mice. For all plots, error bars represent interquartile range.

Supplementary Figure 7

CAR T-cells migration across the BBB-model

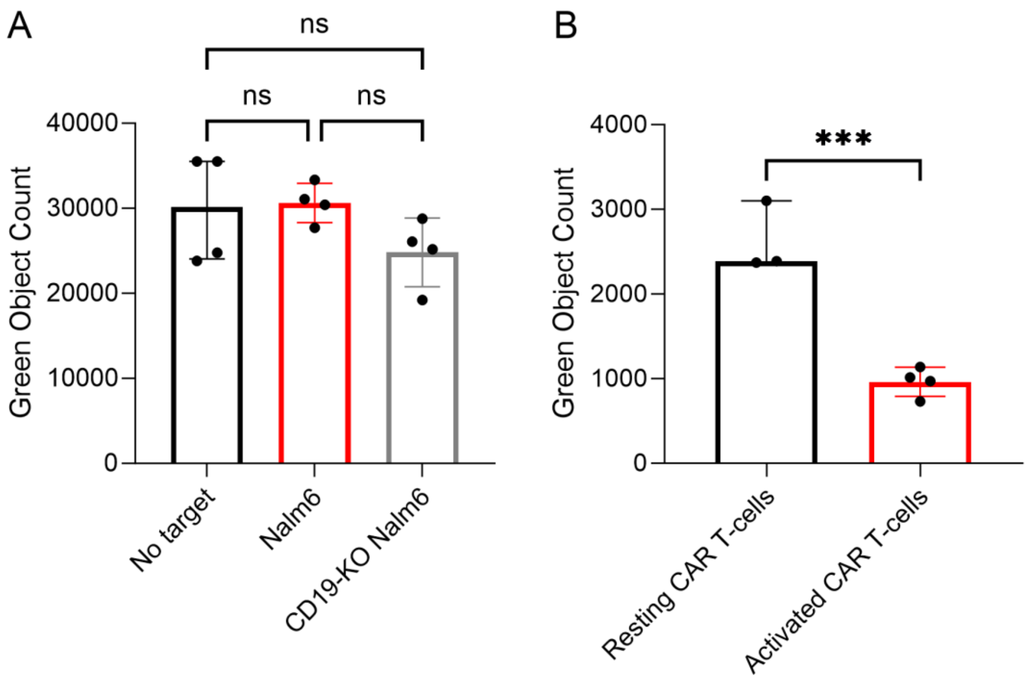


Human CAR T-cells were placed on a human BBB model, consisting of a transwell layered with CD34+ human endothelium and bovine pericytes, and allowed to migrate across the barrier. CAR T-cells were stained with BCECF, and migrated cells were live cell imaged using Incucyte (Essen Bioscience). Analysis was conducted with the Incucyte software (Green object count per image). **A** Migration was performed in the presence of Nalm6 or $CD19^{KO}$-Nalm6 or no target cells seeded in abluminal (brain) compartment of the BBB model. Green object count representing BCECF-stained CAR T-cells in abluminal side are shown. *P* values determined by one-way ANOVA test. **B** Pre-activation of CAR T-cells was performed by co-culture with target cells in a 4:1 effector to target ratio for 24 hours, following by staining CAR T-cells with BCECF and plating them on the luminal side, allowing for 4 hours migration. The green object count represents BCECF stained CAR T-cells in the abluminal side of resting CAR T-cells compared to pre-activated CAR T-cells (*P* value determined by students T test). For all plots, error bars represent interquartile range. $^{ns}p > 0.05$, $^{***}p \leq 0.005$. ‘

Supplementary Figure 8

CAR T-cell exhaustion markers in the BBB-model

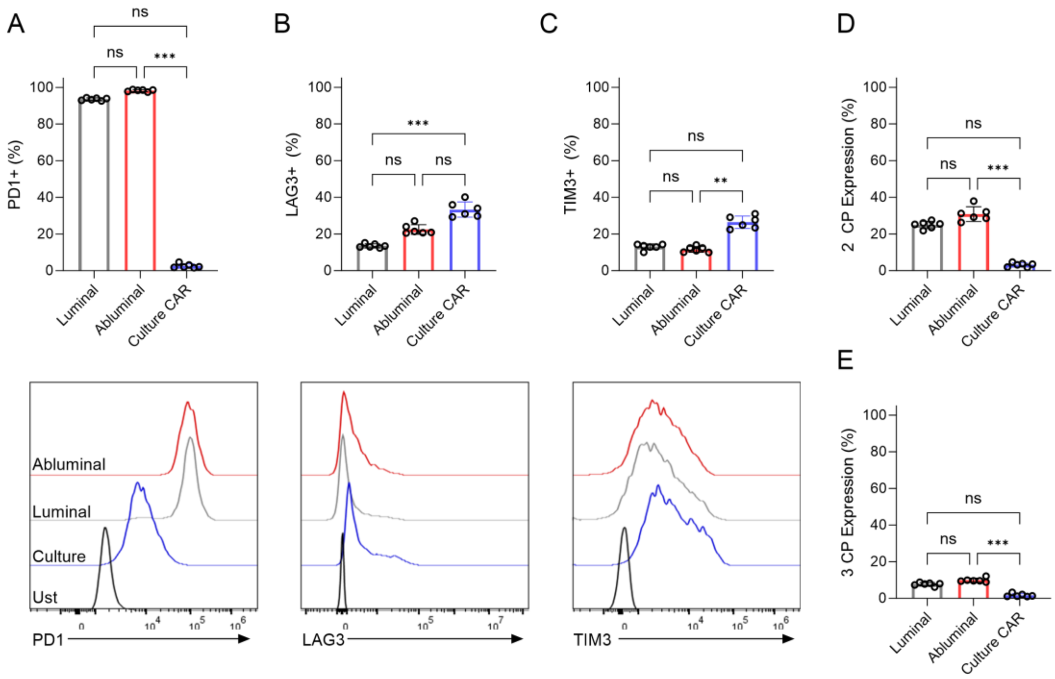


Resting CAR T-cells were allowed to migrate across the BBB model as in Fig. 6. Percentages (upper panel) and representative histograms (bottom panel) of (**A)** PD1+, (**B)** LAG3+, and (**C)** TIM3+ expression on CAR T-cells in the luminal compartment, abluminal compartment and of a BBB-transwell model, as well as the original culture. **D-E** Histograms of the percentage of CAR T-cells expressing 2 checkpoint (**D**) or 3 checkpoint markers (**E**) are shown. For all plots, error bars represent interquartile range. *P* values determined by Kruskal-Wallis test, $^{ns}p > 0.05$, $^{***}p \leq 0.005$.

Supplementary Figure 9

AICD is most prominent when the entire BBB complex is in place

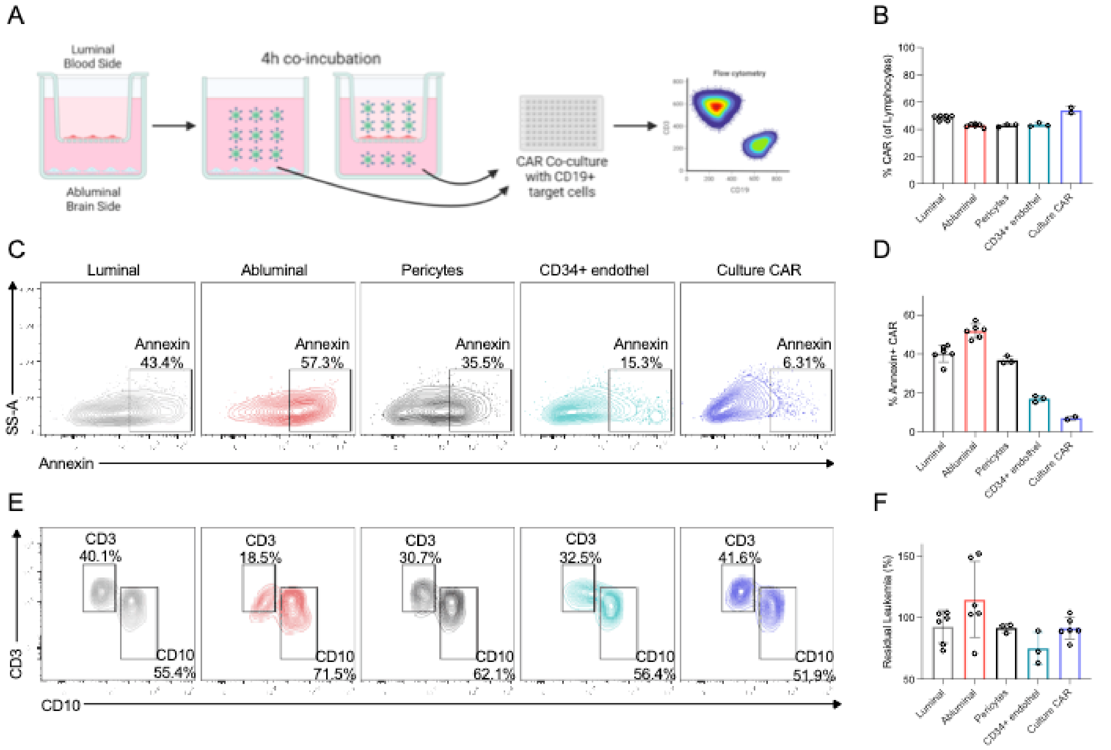


**A** Schematic representation of the human *in vitro* full and separated BBB model migration experiment: CD34+ endothelial cells and pericytes were co-cultured for 6 days in order to enable the endothelium to acquire BBB tight junction. Then 3 transwells (TWs) layered with CD34+ endothelium, were separated from pericytes into a new well containing fresh media. Human CD19 CAR T-cells were introduced to luminal of a full BBB TWs (as in Fig. 6A), to wells containing only pericytes and to TWs containing only CD34+ endothelial cells, and migration was performed for 4 hours. Cells from luminal (grey) and abluminal (red) compartments of full BBB TWs, cells from supernatant of pericytes-only wells (black) and cells from the abluminal side of CD34+endothelium-only TWs (cyan) were collected and either analyzed by flow cytometry, or co-cultured with Nalm6 target cells at a 1:2 effector to target ratio for 24 hours. **B** CAR+ percentage of total cells collected from all conditions, and original culture (blue). **C** Representative flow cytometry plots of Annexin$^{+}$ binding gated on CAR T$^{+}$ cells. **D** Percentage of Annexin$^{+}$-binding CAR T-cells. **E** Representative flow cytometry plots of CD3+ and CD10+ cells after 24h co-culture. **F** Cytotoxicity of CAR T-cells from different compartments was measured by the number of residual CD19-leukemic cells, normalized to target cultured with no effector. For all plots, error bars represent interquartile range.
